# Leveraging ancestral sequence reconstruction for protein representation learning

**DOI:** 10.1101/2023.12.20.572683

**Authors:** D. S. Matthews, M. A. Spence, A. C. Mater, J. Nichols, S. B. Pulsford, M. Sandhu, J. A. Kaczmarski, C. M. Miton, N. Tokuriki, C. J. Jackson

**Author notes:** Authors contributed equally (D. S. Matthews, M. A. Spence).

## Abstract

Protein language models (PLMs) convert amino acid sequences into the numerical representations required to train machine learning (ML) models. Many PLMs are large (>600 M parameters) and trained on a broad span of protein sequence space. However, these models have limitations in terms of predictive accuracy and computational cost. Here, we use multiplexed Ancestral Sequence Reconstruction (mASR) to generate small but focused functional protein sequence datasets for PLM training. Compared to large PLMs, this local ancestral sequence embedding (LASE) produces representations 10-fold faster and with higher predictive accuracy. We show that due to the evolutionary nature of the ASR data, LASE produces smoother fitness landscapes in which protein variants that are closer in fitness value become numerically closer in representation space. This work contributes to the implementation of ML-based protein design in real-world settings, where data is sparse and computational resources are limited.

---

Protein representation learning has been transformative for protein engineering and evolutionary inference^1–7^. Representation learning aims to transform discrete protein sequences, which are inherently high-dimensional and information-sparse, into dense vector representations that numerically capture the salient features of the protein sequence^8^. These representations include the complex relationships between amino acids and provide insights into the functional characteristics and evolutionary histories of the proteins that are embedded in the representation model^3,4,9^. Representations are often provided by protein language models (PLMs) that are trained on large databases of protein sequences^2–4,10–14^. To build representations that capture a protein’s biophysical and evolutionary features in the absence of a functional label, PLMs are often trained by unsupervised masked language modeling (MLM), in which the model is tasked with predicting the identities of residues that have been masked from the surrounding sequence context. The hidden states of a trained PLM, which are retrieved and pooled into a representation, implicitly capture the relevant physical and biological properties of protein sequences^3^. The MLM task itself has also demonstrated utility in protein engineering and evolutionary inference^9,15^. Protein representations can also be derived empirically, such as from the orthogonal principal components of physical amino acid descriptors (i.e. hydrophobicity, solvent accessible surface area, charge)^16–18^; however, such linear representations fail to capture context-dependence, as in deep representation models. Deep representation models, such as PLMs, therefore possess greater context-awareness, which greatly improves their utility in capturing complex biological phenomena.

Protein representation models project sequences to a semantically-rich embedding space that supervised learning models can leverage for supervised tasks (e.g. fitness prediction, where fitness is a quantitative trait of interest, such as catalytic efficiency)^7,19^. This overcomes the inherent challenges of supervised learning in an information-sparse discrete sequence domain that can be confounded by epistasis (i.e. non-linearity in the fitness landscape)^20^. The phenomenon of epistasis is synonymous with ruggedness in the fitness landscape: a highly epistatic system is one in which the effect of a mutation is highly contingent on the background into which it is introduced^21^. In rugged fitness landscapes, a small number of mutations can lead to large nonlinear changes in fitness, making prediction of the functional outcomes of mutations challenging. As a result, high-order epistasis renders molecular evolutionary trajectories unpredictable^22^. Recent work has shown that deep neural networks trained with explicit regularization are able to find smooth representations of protein space^23^, and those that account for epistasis coefficients^24^ can significantly improve the utility of representations in downstream supervised tasks. This implies that embedding spaces that are smooth (with respect to the protein fitness) may be more informative than those that are not. In the case of unsupervised PLMs without explicit regularization, any implicit smoothing in the representation space occurs *via* the learning of contextual sequence dependencies from the MLM objective; therefore, training on sequence data that maximizes the model’s predictive comprehension of epistasis may be expected to produce more informative protein representations.

Ancestral sequence reconstruction (ASR) is a statistical method used to infer extinct molecular sequences from the internal nodes of a phylogenetic tree. Sequences generated by ASR are generally functional and often display novel phenotypes or properties that are ideal for protein engineering^25,26^. Because of this, ASR has been used extensively to generate thermostable proteins,^27–29^ explore functional sequence space^30–33^, and to generate protein scaffolds for engineering^34–36^. Indeed, recent work indicated that ASR outperforms state-of-the-art deep neural machines, including large PLMs, at functional protein generation^37^. ASR has also provided insight into molecular evolutionary processes and understanding sequence-function relationships^38–40^: studying the biophysical, chemical, and biological properties of extinct ancestral sequences can reveal the mechanisms by which proteins acquire novel phenotypes and functions, as well as the features, such as epistasis, that either confound or enable them^41^.

In this study, we describe the use of ASR-generated sequences to train family-specific protein representation models. We show that the incorporation of reconstructed sequences produced representations that surpass state-of-the-art performance on downstream supervised tasks. In the sequence domain, we a correlation between the smoothness of the representation space (with respect to observed fitness) and the predictive strength of the representation. Finally, we show that models trained on large datasets of ancestrally reconstructed sequences have learned a more meaningful representation of epistasis than both equivalent models trained on only extant sequence data and larger PLMs.

## Results

### Multiplexed ancestral sequence reconstruction for protein representation learning

Training deep representation models requires large sequence datasets that cover diverse regions of functional sequence space. From a fully resolved and rooted phylogeny with *n* tips, at most *n-1* ancestral sequences can be reconstructed if only the most likely (the *maximum a posteriori*, MAP) sequence is sampled from the posterior probability distribution of each internal node. This is because ASR is conditioned on a prior phylogenetic hypothesis that inherently limits the sequence diversity generated by ASR to the underlying tree structure (and sequence evolution model)^42^. Here, we have used an approach we term multiplexed ASR (mASR), which samples statistically equivalent topologies as priors for ASR to generate large and diverse sequence datasets approach for representation learning. Because the ground-truth phylogeny cannot ever be known with certainty, phylogenetic topologies are reconstructed by heuristic tree-search algorithms that minimize the negative log-likelihood for a particular tree (e.g.^43,44^). By performing numerous and independent tree searches in parallel, and filtering those that are not statistically equivalent by the approximately unbiased (AU) test^45^, we generate a pool of equally valid, yet distinct phylogenies that are used to reconstruct ancestral sequences. The size of the sampled sequence space means that identical tree topologies are seldom returned by random independent tree searches, effectively increasing the number of sequences produced through ASR by a factor equal to the number of tree-search replicates that are accepted by the AU test.

Another limitation of traditional ASR methods for generating functional sequences is that insertions and deletions (indels) and mishandled or ignored by maximum likelihood sequence evolution models. As indel events are often responsible for driving functional diversification^46–49^, we also developed an automated pipeline for the maximum likelihood reconstruction of insertions/deletions (indels) in large ancestral sequence databases produced with mASR (SI Fig 1). For site *n*, all tips are assigned a binary label (0 for no indel, 1 for indel), depending on whether a gap character is observed. We assume that the rate of insertion in a sequence is approximately equal to the rate of deletion and use an equal-rates model to reconstruct the probability of a gap being present at site *n* of all ancestral nodes. We repeat this for all sites where a gap is observed in >= 1% of the extant sequences and remove all residues from ancestral nodes reconstructed with a label of < 0.5. Our maximum likelihood indel processing combined with mASR allowed us to generate large datasets of realistic sequences from evolutionary information.

In this study, we applied mASR with our indel processing pipeline to the bacterial phosphotriesterase (PTE) family (SI Fig 2), a group of homologous proteins that hydrolyse synthetic phosphotriesterase pesticides and that has been used extensively as a model for functional adaptation and molecular evolution^50–53^. From an initial alignment of 293 non-redundant, extant PTE homologues, we generated a dataset of >10000 unique PTE-like sequences with 100 replicates of tree-search, despite most replicates returning highly similar topologies (Fig 1a). To investigate the diversity of sequences generated by mASR, we embedded ancestral and extant sequences in the PLM ESM-1b^3^, which has been pre-trained by MLM on ∼250 million non-redundant protein sequences in the UniRef50-S database. We then used t-distributed stochastic neighbor embedding (tSNE) to project the ESM-1b representations of extant and ancestral PTEs onto 2 dimensions for visualization (Fig 1b). The majority of the ancestral sequences produced with mASR belong to regions of sequence space that are not sampled by the extant PTE homologues used to reconstruct them. This, combined with the fact that ancestrally reconstructed proteins often feature increased thermostability and comparable catalytic activity to the extant sequences used to reconstruct them^25^, means that mASR may serve as a source of sequence novelty. As ancestral sequences are not represented in large, pre-trained PLMs, ancestral-like sequences are highly unlikely to be produced by generative PLMs designed specifically for novel sequence generation in protein engineering, meaning that mASR (including the maximum likelihood indel reconstruction) is a unique statistical method for the generation of large ancestral sequence datasets.

**Fig 1.**
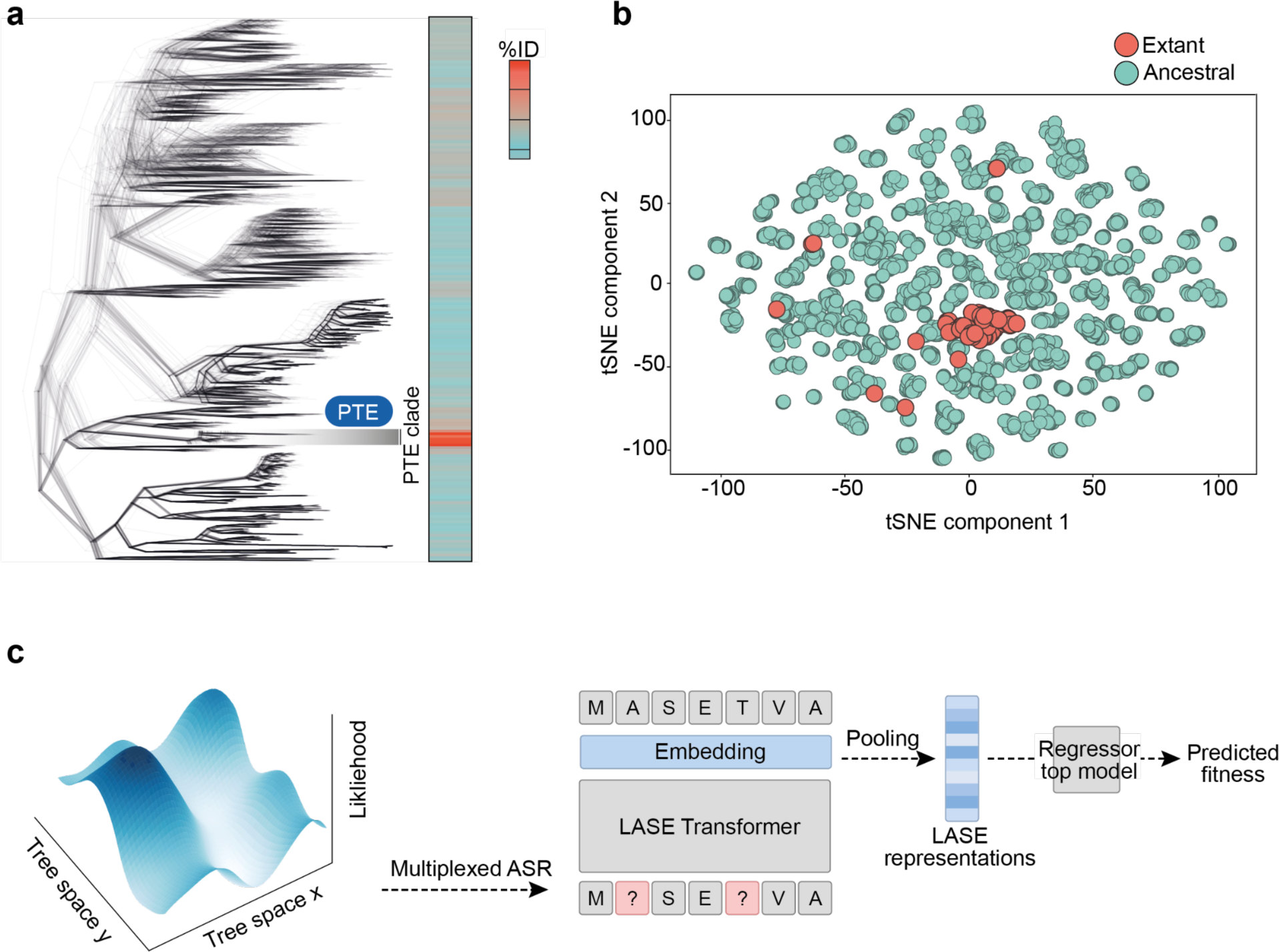
The use of mASR in the LASE pipeline. (a) Using mASR, 100 statistically equivalent phylogenetic trees were produced from an alignment of non-redundant, extant PTE homologues. Similarity to the wt PTE enzyme (shown in blue, uniprot ID: P0A434) is sparse, and limited primarily to few sequences that belong to the lineage that wt PTE diverged from. (b) The tSNE projection of ESM-1b embeddings of the ancestral sequences obtained from mASR (cyan) as well as extant sequences (salmon). MASR increases the sequence volume and diversity of PTE-like enzymes beyond natural diversity. (c) LASE diagram. Datasets of ancestrally reconstructed sequences produced by mASR are used for MLM in a small transformer. Residue-wise embeddings are pooled into fixed length representations, which are used in downstream supervised learning with a regressor top model.

#### Local ancestral sequence embedding (LASE)

We used the datasets generated by mASR to train small transformer encoders for representation learning (Fig 1c)^54^. Transformers consisted of a positional embedding layer, 6 encoder blocks, each with a 4-headed multi-head attention (MHA) layer and a feed-forward fully-connected layer, and a time-distributed fully-connected output layer. Embedding and feed-forward dimensions were 128 and 512, respectively, and all models were trained using an MLM objective. The encoder blocks of the transformer learn long-range sequence dependencies in the training data from the MLM objective; the hidden states of these layers capture the physical, chemical, and biological properties of the protein sequence being embedded and are pooled into dense, fixed-length protein representations, as with previous transformer-based PLMs^2,3^. The small size (∼2.3 M parameters) of these representation models means they are computationally inexpensive to train, requiring <1 hour per epoch on conventional hardware (single Nvidia A10). This is in contrast to large PLM fine-tuning (SI Fig 3), which requires retraining of all parameters on family or task-specific data^3,4^, or spiked data scheduling, where the fine-tuning dataset size is diluted 100-fold with unrelated protein sequences to avoid forgetting of universal protein features^5^. We term this approach local ancestral sequence embedding (LASE) as it uses synthetic ancestral sequence data to build an informative representation space for a local sequence space sampled through mASR.

We then tested the predictive performance of LASE. The arylesterase activity of PTE variants for 2-naphthyl hexanoate (2NH) has been previously described in four independent experiments. Three of these are directed evolution studies: two “forward” evolution experiments in which arylesterase activity was selected for in two independent experiments from the wt enzyme (R-trajectory^55^ and S-trajectory^56^) and a “reverse” evolution experiment in which the arylesterase obrainted at the end of the R-trajectory directed evolutuion experiment was evolved to regain PTE activity^57^. The fourth dataset originates from to a study in which the combinatorial mutagenesis of residues near the wt PTE active site was used to investigate epistasis^58,59^. Similar approaches have recently been developed and used to study the fitness landscape of the enzyme dihydrofolate reductase^60^. To produce a standard training dataset, data from each experiment were combined by normalizing activity measures around data points common to all experiments (SI 2). The structure of the final dataset is depicted as a Hamming graph (Fig 2a) where two distinct data structures are seen: trajectory data, where mutations are accumulated along linear directed evolution trajectories, and combinatorial data, where all combinations of 6 active site substitutions are explored.

**Figure 2.**
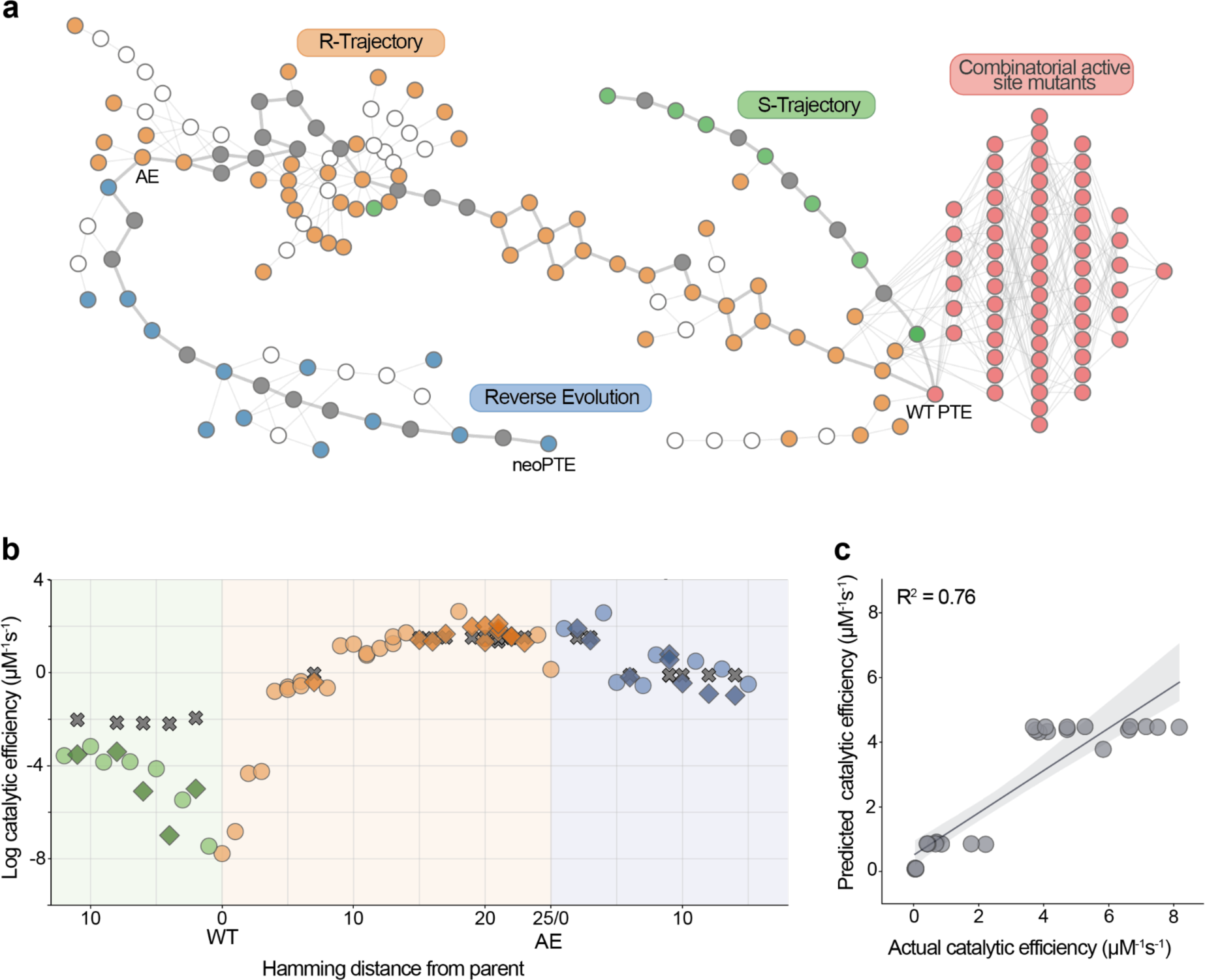
LASE results on the PTE test system. (a) The PTE dataset depicted as a graph, where nodes represent unique sequences and edges represent one-Hamming distance (one mutation). Nodes are colored by their experimental origin: R-trajectory (orange)^55^, S-Trajectory (green)^56^, reverse evolution (blue)^57^, combinatorial active site (salmon)^58,59^, sequences tested in this study (grey). Data points seen in white have not been experimentally characterizsed. (b) PTE variants from the trajectory data are visualized by log catalytic efficiency against Hamming distance from the parent sequence that evolution was initiated from. Circles represent training data from trajectory datasets, diamonds represent the experimentally determined values for the test dataset, and crosses represent catalytic efficiency predictions from the best LASE random forest (RF) regressor model. (c) The performance of the best LASE RF regressor on test data as the predicted catalytic efficiency against the observed catalytic efficiency.

LASE representations were benchmarked against one-hot encodings (OHE), learned representations from PLMs trained on large databases (UniRep^4^, ProtTrans^10^, ESM-1b^3^ and ESM-2^2^), as well as empirical representations derived from the physicochemical properties of amino acids (Georgiev^16^, Z-scale^17^, ProtFP^18^). To assess the performance of LASE, we synthesized and assayed 26 PTE mutants that were not sampled in directed evolution trajectories (grey nodes, Fig 2a), and compared the predicted and observed arylesterase catalytic efficiencies. Each of these sequences carried the intermediate single mutation between directed evolution generations *n* and *n+1*, for all generations separated by more than a single mutation. A range of model architectures were trialed for the supervised regression task (SI Table 1; SI Table 2). The performance of each representation model was measured as the Pearson’s R^2^ correlation coefficient between the observed and predicted fitnesses in this test set. This metric is a valid reflection of real-world predictive performance as the data from train and test sets were produced from independent experiments, as is often the case in protein engineering workflows.

When trained on a 0.5% masking objective, LASE was the most informative representation for predicting arylesterase catalytic efficiency, scoring a Pearsons’s R^2^ of 0.76 over the test data. Current state-of-the-art ESM representation models, ESM-1b and ESM-2, performed significantly worse than LASE at catalytic efficiency prediction, achieving Pearson’s R^2^ scores of 0.69 and 0.62 respectively, despite having >300-fold more parameters than the LASE representation (670 M parameters, trained on 250 M sequences for ESM-1b *vs.* 2.3 M parameters, trained on 10000 sequences for LASE). Interestingly, there is a significant difference in the performance of representations from LASE models trained on 0.5% and 15% masking objectives (Pearson’s R^2^=0.76 and 0.54, respectively) (SI Figure 6), indicating that masking percentage is a hyperparameter that should be optimized during PLM training. Indeed, masking 0.5% of the PTE sequence (∼2 sites per training sequence) may intuitively provide representations that capture richer information on the scale of the test dataset (typically single mutational differences) than the conventional 15% masking (∼54 sites per training sequence) objective.

The sequences of variants in the PTE dataset were embedded using each representation scheme and a random foresrt (RF) model was optimized (3-fold cross validated with 10 randomized 20% evaluation holdout sets). The final models were trained on all available data and performance was measured on the true holdout set of 26 variants from within the trajectory data (interpolation set).

To test whether the performance of our approach could be replicated without using ancestrally reconstructed sequences in MLM training, we next trained a model identical to the best performing LASE model on only extant PTE homologues. In other words, we tested whether the performance was a result of the evolutionary nature of the sequence data. To ensure fair comparison between the models, the training task (0.5% masking percentage, trained for 25 epochs with an early stopping patience of 2 epochs) and the number of training tokens that either model was exposed to during MLM were kept constant. Representations from the extant-only model (local extant sequence embedding; LExSE) were significantly less informative than representations from the equivalent LASE model, scoring a Pearson’s R^2^ of 0.56 (Table 1), indicating that ancestrally reconstructed sequences are fundamentally more informative for training representation models than homologous extant sequences are. It has similarly been observed that incorporating evolutionary inference to representation learning improves predictive performance on downstream tasks^61,62^.

**Table 1.** Benchmarking of representation schemes on the PTE dataset.

| Representation | Interpolation set performance ($R^2$) |
| --- | --- |
| One-hot | 0.60 |
| Z-Scales | 0.57 |
| ProtFP | 0.58 |
| Georgiev | 0.59 |
| UniRep | 0.56 |
| ProtTrans | 0.63 |
| ESM-1b | 0.69 |
| ESM-2 | 0.62 |
| **LASE** | **0.76** |
| LExSE | 0.56 |

### Computationally efficient *in silico* evolution

Due to the inherently small size our models, we next investigated the improvement in inference speed our approach provides relative to current state-of-the-art representation models. We performed *in silico* evolution in the discrete sequence domain to quantify the wall-time acceleration that LASE provides, relative to the next best performing representation model, ESM-1b ^3^.

We performed evolution through generations of diversification and selection. During diversification, all single mutations in the post-selection pool of sequences at generation *n-1* are recombined into the pre-selection pool of generation *n*. This pool of sequences is embedded in the representation model for catalytic efficiency prediction. During selection, the 250 highest ranked sequences from the generation *n* pre-selection pool are carried forward to the next generation of diversification. To avoid becoming trapped in local optima, we also allowed an additional 250 randomly sampled sequences to continue through each generation of evolution. On conventional consumer hardware (a single Nvidia RTX 3080) without background processes running, we observe an overall >75-fold reduction in the wall-time required for sequence embedding for full generational inference on a per-sequence basis between LASE (∼22 seconds per 40000 sequences) and ESM-1b (∼26 minutes per 40000 sequences) (Fig 3a). Over 25 generations of *in silico* evolution, LASE-based evolution converges at the predicted fitness peak within 2 minutes, whereas the equivalent ESM-1b-based evolution converges at an equivalent predicted fitness peak after 2 hours on identical hardware (Fig 3b). Recent work on protein G and immunoglobulin G has indicated that neural machines can accurately extrapolate beyond the sequence space of the training data^63^, suggesting that this approach *in silico* evolution may sample the PTE fitness peaks not yet discovered by directed evolution.

**Fig 3.**
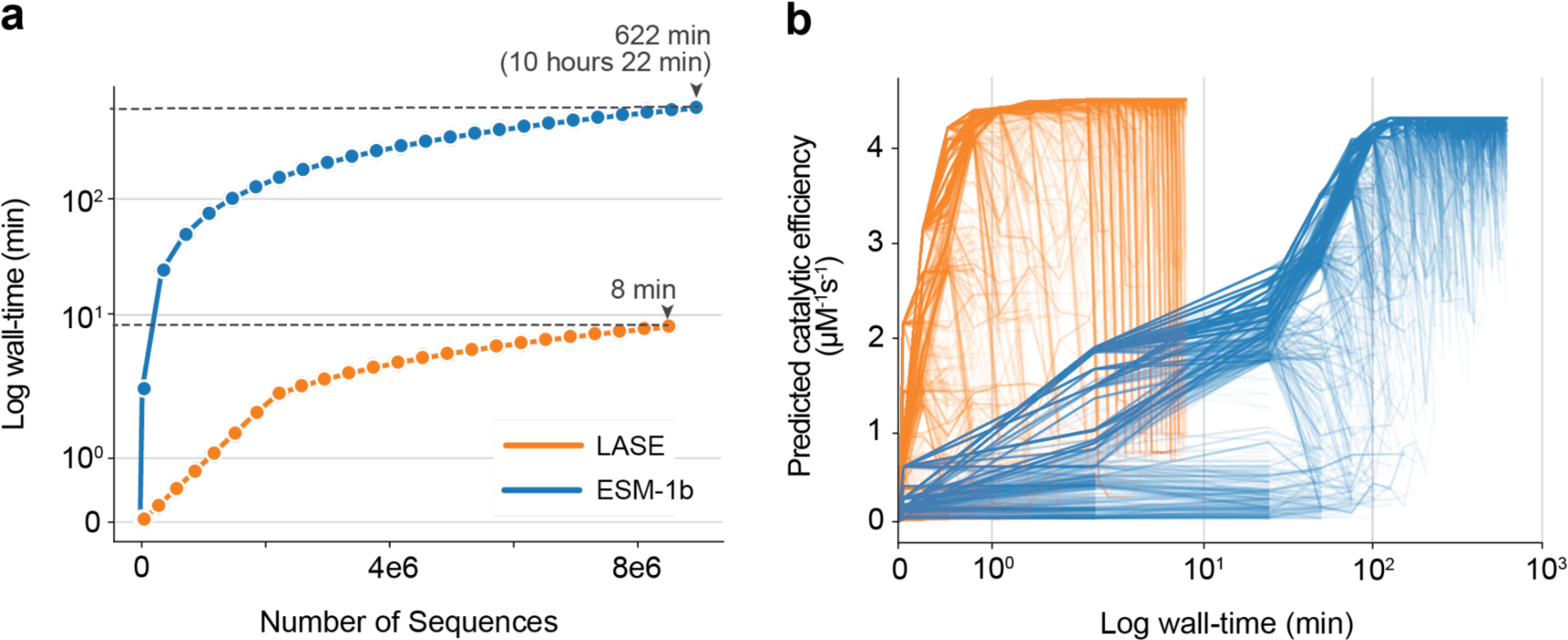
LASE and ESM-1b embedding wall-time. (a) Embedding time as a function of the number of sequences to be embedded for LASE (orange) and ESM-1b (blue). LASE provides a >75-fold decreases in the required wall-time. (b) Predicted catalytic efficiency as a function of wall-time for LASE (orange) and ESM-1b (blue) *in silico* sequence evolution. All evolutionary trajectories for LASE converge at the fitness peak within 1.2 minutes, whereas ESM-1b-based optimization requires more than 2 hours to converge.

#### Smooth representation space underlies predictive performance

Having established that LASE representations surpass the performance of models trained with large numbers of extant sequences on supervised tasks for PTE, we next sought to discern the relationship between the structure of each model’s representation space and predictive performance. We hypothesized that the smoother an observed fitness is on a model’s embedding space, the better the predictive performance of that model’s representations may be on a downstream supervised task using a regression top model. This hypothesis is based on the premise that representations that embed sequences in such a way as to make the fitness function smooth require a less complex transformation to the fitness domain than representations yielding rugged representation spaces, where there are complex relationships between the representation vector and the fitness. Indeed, it has been previously shown that explicitly smoothing the representation space of a transformer with respect to fitness results in more informative representations on downstream regression tasks ^23^.

We took a spectral graph approach to assess how rugged/smooth the observed fitness is under each model’s representation embedding for the PTE dataset^64^. This involved using the normalized Dirichlet energy^64–66^ as a measure for smoothness on *k*-nearest neighbor (KNN) graphs of the embedded PTE dataset, where nodes represent embedded sequence coordinates and edges connect the *k*-closest neighbors between each embedded sequence (SI Fig 4). In general, Dirichlet energy is an effective measure of how rugged (non-smooth) a function is^64^. Here, the local Dirichlet energy is the squared difference in observed fitnesses between adjacent nodes in the KNN graph; in KNNs where nearby nodes map to similar numerical fitness values, the Dirichlet energy is low, and the embedded fitness landscape can be said to be smooth.

Across all tested representations, we find a negative relationship between the normalized Dirichlet energy of representations’ embedded KNN graphs and the RF regression model’s predictive performance (Pearson’s R^2^ correlation coefficient)(R2 = 0.67, P=0.0016) (Fig 4a; b). This indicates that the smoothness of the representation spaces (with respect to the fitness) generally correlates with more informative representations in the PTE dataset. LASE yields the smoothest embedded KNN-graph, consistent with performance on arylesterase activity predictions on the PTE test set (Table 1). When these representations are projected onto a 2-dimensional tSNE basis for visualization, ESM-1b projections resemble the underlying Hamming graph structure in the sense that the S-Trajectory PTE variants with high arylesterase catalytic efficiencies are projected separately from R-Trajectory PTE variants with high arylesterase activity (Fig 4c). In contrast, projections of the LASE embedding space (Fig 4d) group these functional variants as neighbors (i.e. they are clustered by fitness), despite their disparate evolutionary history.

**Fig 4.**
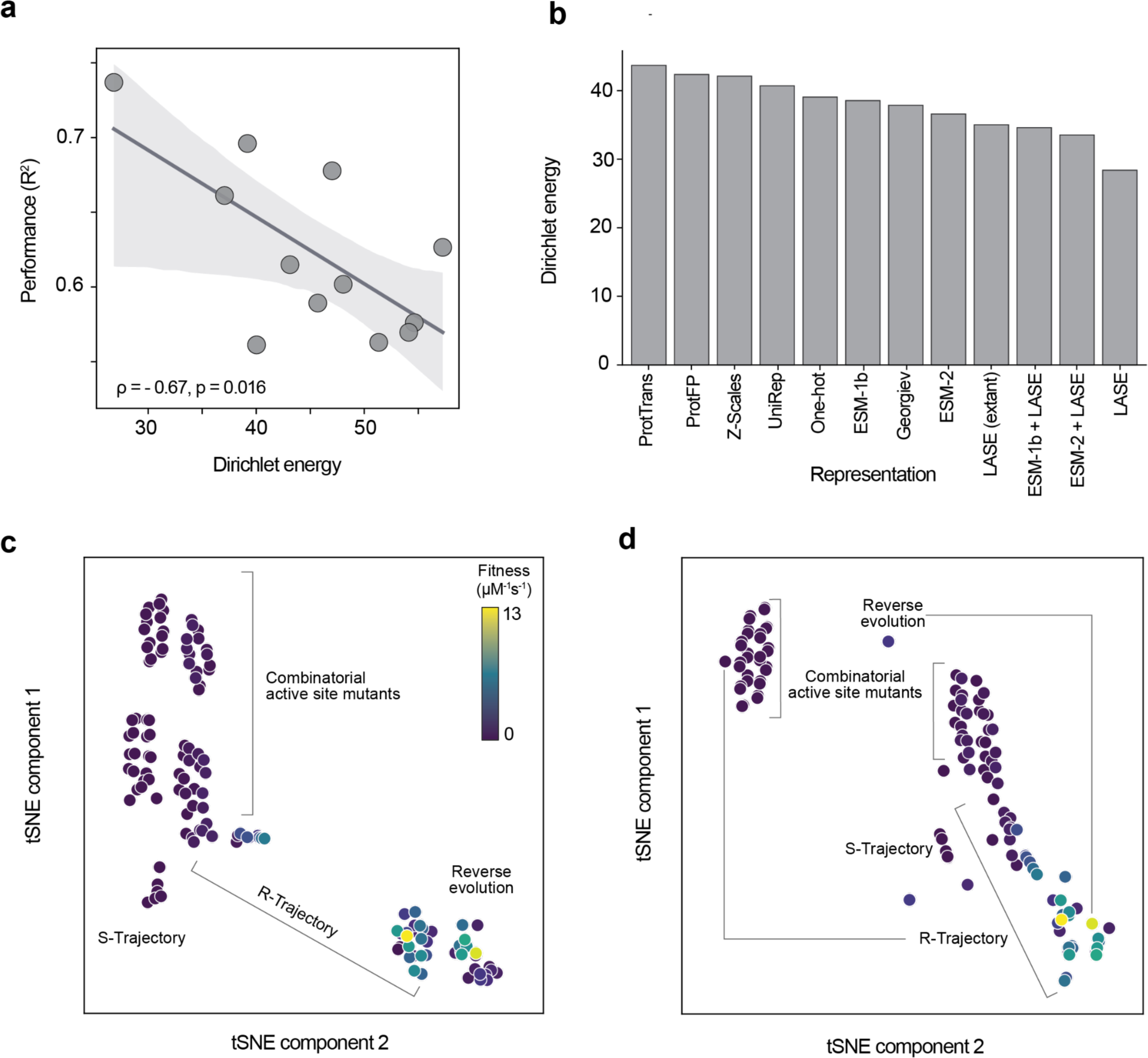
Comparison of representation space structure. (a) Correlation between the Dirichlet energy of a *KNN* graph embedded in a model’s representation space and the predictive performance (R^2^) of that model. There is a significant correlation between the smoothness of a representation space and the performance of the representation model (p=0.016). (b) The normalized Dirichlet energies for each representation scheme. (c) The ESM-1b and (d) LASE representations projected into a 2-dimensional space with tSNE and colored by arylesterase catalytic efficiency. LASE clusters highly functional variants together, independently of their evolutionary history.

#### Ancestrally reconstructed sequences are more informative for learning epistasis than extant sequences

We next compared the topological features of LASE and the equivalent local extant sequence embedding (LExSE) embedded PTE fitness maps to assess whether ancestrally reconstructed sequences are fundamentally more informative in learning epistasis than their extant counterparts. To do this, we first computed the Dirichlet energy of each node’s immediate neighborhood from the embedded KNN graph. If the energy of a node-wise subgraph is high, the fitness of that node deviates in an unpredictable way from its immediate neighbors and the local representation space is rugged with respect to fitness. We complemented this with graph spectral decomposition. Whereas the node-wise Dirichlet energy is a metric of how ‘strained’ a node is given its context in the graph, spectral decomposition provides insight on the contributions that low frequency eigenmodes from the graph topology make towards the observed fitness landscape. When the fitness landscape over a graph is smooth, it is decomposed into low frequency (i.e. simple) eigenmodes with greater magnitudes than high frequency (i.e. complex) eigenmodes^67–69^.

We systematically observe lower, and hence less ‘strained’, local Dirichlet energies in the LASE representation space than the LExSE representation spaces (Fig 5a - c). Interestingly, while the LASE KNN graph is less strained than other representations, local energies in each graph follow the same general node-wise structure, indicating that the sequences most confounded by epistasis are common between representations and embedding in a representation model only dampens complexity in the fitness map, rather than drastically altering it. In both representation spaces, for example, the node with the greatest Dirichlet energy belongs to the qudaruple mutant D233E/H254R/L271F/F306I, which is the fitness peak in the combinatorial fitness map (Fig 5a). This result is corroborated by spectral decomposition, where lower frequency eigenmodes over the graph topology make greater contributions to the fitness function in the LASE space than they do in the LExSE space (and vice versa for high order eigenmodes; Fig 5d). Finally, when the PTE combinatorial fitness map is projected from the LExSE and the LASE representation spaces to two dimensions with tSNE for visualization, we observe a clear functional gradient along both components of the LASE space that are absent in the corresponding LExSE space (Figure 5e, f). Moreso, functional clustering is not the consequence of the underlying dataset structure as sequence clusters in the tSNE basis are not arranged by the number of mutations from the wt sequence (Fig 5g). These results together demonstrate that smoothness in the representation space of LASE is a consequence of learning the MLM task on ancestrally reconstructed sequences and cannot be produced with comparable extant-only sequence data.

**Figure 5.**
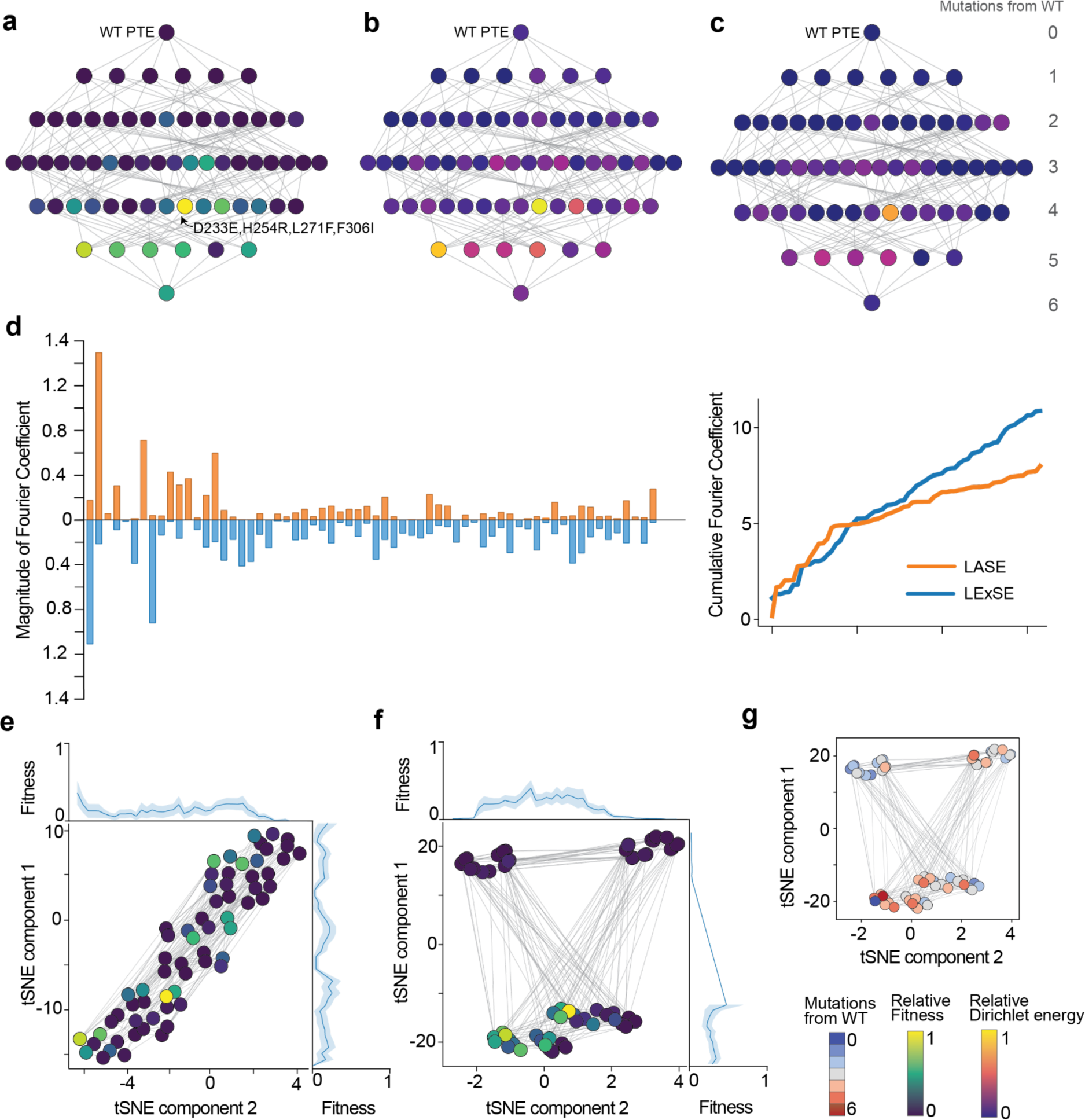
ASR sequences allow for the learning of epistasis. Local ruggedness in the PTE combinatorial fitness map. (a) Hamming structure of the combinatorial PTE active site. Nodes represent unique sequences and edges connect nodes separated by a single mutation. Nodes are coloured by arylesterase catalytic efficiency. The fitness peak (D233E/H254R/L271F/F306I) is labeled. (b) Node-wise (local) Dirichlet energies of the combinatorial PTE fitness map when embedded in LExSE and (c) LASE. Nodes are coloured by Dirichlet energy determined over subgraphs that include a node’s immediate neighbors. Local ruggedness is higher in the LExSE embedding space than the LASE embedding space. (d) Graph Fourier transform of LExSE (blue) and LASE (orange) KNN graphs. Higher magnitudes at lower eigenvectors (ordered along x-axis) indicate greater contributions from low order (i.e. non-complex) eigenmodes, and hence smoothness in the fitness landscape. (e) PTE combinatorial dataset embedded in LExSE and (f) LASE. Nodes are coloured by catalytic effiency and average fitness is projected to axes. (g) LASE embedded KNN graph where nodes are coloured by distance from the wt sequence. Structure in the embedding space of LASE is not a consequence of the underlying data structure (i.e. (a)).

## Discussion

There has been significant recent development in generative deep learning models to produce diverse, functionally-enriched sequence datasets^11,12,70–72^. Here we have presented mASR, which achieves this by leveraging phylogenetic information. Unlike mASR, deep generative models memorize the features of protein sequence space in the weights of a neural machine architecture. This is an intensive task as it often requires regressing (e.g.^11^) the predicted tokens as sites *<n* to predict the identity of the token at site *n,* where *n* is often hundreds of tokens in length and output sequences are remarkably diverse. ASR alternatively leverages the sequence information that is jointly encoded in a phylogenetic hypothesis, removing the implicit requirement to memorize sequence features within the model. While the model we have trained is not holistic in that different protein families require independent ancestral sequence reconstructions, it is not dissimilar from state-of-the-art deep generative models that typically require fine-tuning on a per sequence family basis for accurate sequence generation^11^. Moreso, systematic biases that lead ASR to produce thermostable yet functionally diverse proteins^25^ make mASR a compelling approach to novel sequence generation when compared with deep generative alternatives^37^.

The predictive performance of a representation model is improved when evolutionary information from mASR is incorporated into MLM training. Here we have shown that embeddings from LASE and LExSE differ significantly both in their predictive capacity and their comprehension of epistasis. These results indicate that ancestrally reconstructed sequences are inherently more informative for representation learning than extant sequences. One intriguing hypothesis is that the representation model learns meaningful differences between sequences by building an implicit understanding of their genealogical relationships. Indeed, the observation that identical mutations have different phenotypic effects across homologous protein backgrounds suggests that their patterns of epistasis have diverged from one another^31,73–76^. With only extant sequence information, however, features of sequence divergence are ambiguous. By learning the MLM task on ancestrally reconstructed sequences as they diverge under different selective pressures, it is reasonable to hypothesize that the embedding model will learn representations that capture epistasis, as well as the sequence features responsible for functional divergence.

A key insight from this work is the role ruggedness in an embedding space plays in that embedding space’s utility. We demonstrate that the smoother a model’s latent space is with respect to fitness, the more interpretable, and hence informative, that space is in the PTE system. One hypothesis for how PLMs achieve this is through re-arranging the discrete sequence domain into a numerical domain informed by protein structure, evolutionary relationships, amino acid properties and long-range dependencies (i.e. epistasis), which are implicitly captured in the sequence and each relate to the protein’s function^2,3,5,8^. This latter embedding represents proteins’ relevant features more faithfully, yielding a smoother (i.e. less complex) function from the embedding space to the fitness values that can be more readily leveraged by downstream supervised models. While our approach is unique in achieving this through ASR, we are not the first to make the observation that smoother representation spaces in protein representation models are more interpretable, and hence informative for downstream supervised tasks^23^. Unlike other methods, however, our approach does not consume experimental data during model training for explicit latent space smoothing.

Here, we have used small, family-specific transformer representation models to emphasize the computational efficiency of our approach, however an interesting direction may be large PLM fine-tuning on ancestrally reconstructed datasets. While we have demonstrated that ancestrally reconstructed datasets produce topologically more informative latent spaces than extant sequences, it is unclear to what extent (if at all) this may transfer to large PLM fine-tuning. Similarly, it is unclear to what extent more explicit evolutionary information may improve the usefulness of PLM representations. Indeed, our approach ignores the inherently sequential nature of ASR over a phylogenetic tree and instead allows the model to learn these features implicitly through the MLM task. A training task such as next sequence prediction that is analogous to the next sentence prediction training task of BERT^77^, for example, may fully and explicitly leverage the evolutionary information encoded in ASR for representation learning.

## Methods

### Multiplexed ASR

Using wt PTE (uniprot ID: P0A434) as the query, 1000 sequences with homology to PTE were retrieved from the NCBI NR database by BLAST. Redundancy in this dataset was removed to 90% sequence identity using CD-HIT^78^. A sequence alignment was generated using the GINSI protocol of MAFFT^79^, which was manually refined by removing sequences that aligned poorly, or featured gaps in conserved regions of the protein. 100 independent replicates of tree-search were performed in IQ-TREE 2^43^, with complete model parameterization by ModelFinder^80^ (as implemented in IQ-TREE 2). The AU-test^45^ was conducted to 10000 replicates on all 100 converged tree topologies. Ancestral states were reconstructed in IQ-TREE 2, ancestral sequences were extracted as the MAP character at each site of the input alignment.

To process indel events by maximum likelihood in ancestral sequences as part of the mASR pipeline, each extant sequence in the alignment was attributed a binary vector with length equal to the number of insertions in the alignment, where 0 at dimension *i* denotes a gap character at alignment index *i* and 1 at dimension *j* denotes a non-gap character at alignment index *j*. Insertions are only considered for sites where ≥1% of residues feature a gap character. Using a Jukes-Cantor-like model that assumes the rate of insertion and deletion are equal (e.g. the rate of 0 transitioning to 1 is symmetrical), we reconstruct the ancestral character 0 or 1 (e.g. insertion of no insertion) by maximum likelihood as a discrete trait for node *n* at site *i* for all nodes and sites in the tree and insertion sites of the alignment, respectively. The ancestrally reconstructed character at site *i* of node *n* remains if the corresponding discrete trait is reconstructed with a likelihood of ≥ 0.5, or is otherwise replaced by a gap character. Maximum likelihood discrete trait reconstruction was performed using the ACE function from the APE package in R^81^.

### LASE

LASE models were implemented in TensorFlow 2.13.0. Models consisted of a 128-dimensional sinusoidal positional embedding layer, six encoder blocks with 4-headed multi-head attention (128 dimension) and a feed-forward fully-connected layer (512 dimension)^54^. The hidden states from the last encoder block were passed to a time-distributed fully-connected layers with 20 output neurons with softmax acitvations. Loss was determined as the categorical cross entropy loss, as natively implemented in TensorFlow 2.13.0. Each model was trained for 25 epochs with an early stopping criteria of 2 epochs without decrease in the validation loss using the Adam optimizer^82^ (as implemented in TensorFlow 2.13.0) with the default learning rate of 1e-3. Input sequences were tokenized and pre-masked with 0.05% and 15% masking probabilities. Models were trained with a batch-size of 128.

### Data Normalization

Labeled PTE datapoints comprising sequence and catalytic efficiencies were obtained and normalized from four sources: three directed evolution experiments as well as an active site study^55–59^. This was necessary as all training data were retrieved from different studies with slight differences in experimental protocols. Datapoints where the variant activity was labeled as being relative to another variant were converted to catalytic efficiency (Supplementary Information 2), then data from separate experiments was normalized through linear scaling. Linear scaling was applied to local regions of experiments where possible. For instance, when possible, variants were normalized with respect to a single neighboring variant for which the catalytic efficiency was known, rather than all variants in the experiment where the catalytic efficiency had been measured.

### Regression Models

Regression models were implemented with scikit-learn 1.2.1. Prior to modeling, all fitness labels were scaled using min-max normalization. Hyperparameter optimization was conducted through Bayesian Optimization with scikit-optimize 0.9.0 (Supplementary Information 3). Model architectures tested and their corresponding hyperparameter search space are listed in SI Table 1. For the training dataset (n=136), a 20% holdout set was used for testing and hyperparameter optimization was performed over a 3-fold cross-validation data split. This was repeated over eleven randomized holdout sets (each with different random-states) to determine average model performance. Representation extraction methods are outlined in Supplementary Information 4.

### In silico evolution

Evolution commenced on the wt PTE variant and ran for 25 generations of mutation and selection. To reduce the size of the explored sequence space to a manageable size, the aligned mASR data was used to determine which mutations may arise at each site of the sequence. Specifically, residues that existed in at least 10% of the aligned data were considered to be a “plausible” mutation. Further, no insertions or deletions were allowed. For each generation, all possible mutations of the previous generation were made, the catalytic efficiency was predicted, the 250 variants with the best catalytic activity as well as an additional 250 random variants were selected as the final generation and all variants that had previously been generated were removed.

### Dirichlet Energy Calculations

To estimate the ruggedness of the combinatorial PTE landscape, for different representation schemes, k-nearest neighbor (KNN) graphs with *k* = √*N* were produced with scikit-learn 1.2.1. The kNN graphs were made symmetric by allowing for an edge in either direction to be a single edge. The dirichlet energy was calculated as previously described^66^ and is defined as:

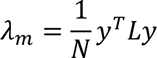

Here, *λ_m_* represents the normalized dirichlet energy, *N*, the number of nodes in the graph, *y*, the signals (fitness values) of each node and *L* the graph Laplacian operator of the adjacency matrix of the kNN graph.

### Fourier Spectral Decomposition

The empirical Fourier coefficients were determined using kNN graphs of the sequence space. First, the generalized eigenvalue problem on the graph Laplacian was solved using the ‘eigh’ function from the Python package SciPy to determine eigenvalues and eigenvectors for the space. The Fourier coefficients were then determined from the eigenvectors as follows^83^:

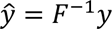

Where, 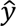 are the Fourier coefficients (graph Fourier transform), and *F*^−1^ are the eigenvectors.

The absolute values of 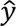 were then reported as the eigenmodes at frequencies that correspond to the ranked eigenvalues.

### Protein expression

Genes encoding the 26 PTE variants in the test set were ordered from Twist Bioscience (cloned into the pET29b plasmid with a C-terminal hexahistidine tag) and transformed into *E. coli* BL21(DE3) cells through electroporation. The *E. coli* was plated onto lysogen broth (LB) supplemented with 500 μM Kanamycin and grown overnight at 37°C. Starter cultures were grown in 1mL of LB media supplemented with 1mM Kanamycin with shaking at 37°C to OD_600_ = 0.6. 5mL of 37°C LB media supplemented with 1mM Kanamycin was then inoculated with approximately 2e8 *E. coli* cells and grown at 37°C with shaking to OD_600_ = 0.6. Protein expression was induced with 1mM IPTG. After 1 hour of growth at 37°C, cells were harvested through centrifugation at 3300 rpm for 10 minutes at 4°C. The resulting pellet was then stored at −80°C before being resuspended and lysed in 200 μL Lysis Buffer (50 mM Tris-HCl, 1x BugBuster Protein Extraction Reagent, pH 7.5) for 30 minutes at room temperature. The final protein lysate was then collected via centrifugation at 3320 rpm for 30 minutes at 4°C. Activity was measured from the cell lysate.

### PTE characterization

For PTE characterization, a 96-well plate assay was conducted. Reactions were performed in transparent 96-well plates containing 20 μL of clarified cell lysate and 180 μL of 200 μM 2NH (Sinfoo Biotech) in 50 mM Tris-HCl, pH 7.5 supplemented with 0.1% Triton-X100. 2NH hydrolysis was monitored at 500 nM through complex formation with Fast Red (SISCO). Initial rates of formation were normalized to cell density (OD_600_) and average values were used as final relative activity measures. Relative activity values were then normalized by linear normalization of data to previously measured PTE variants^55,57,58^ that were also expressed and characterized.

## Availability

The code for model training and analysis is available on GitHub https://github.com/RSCJacksonLab/local-ancestral-sequence-embeddings.

## Supplementary Information

**SI Fig 1.**
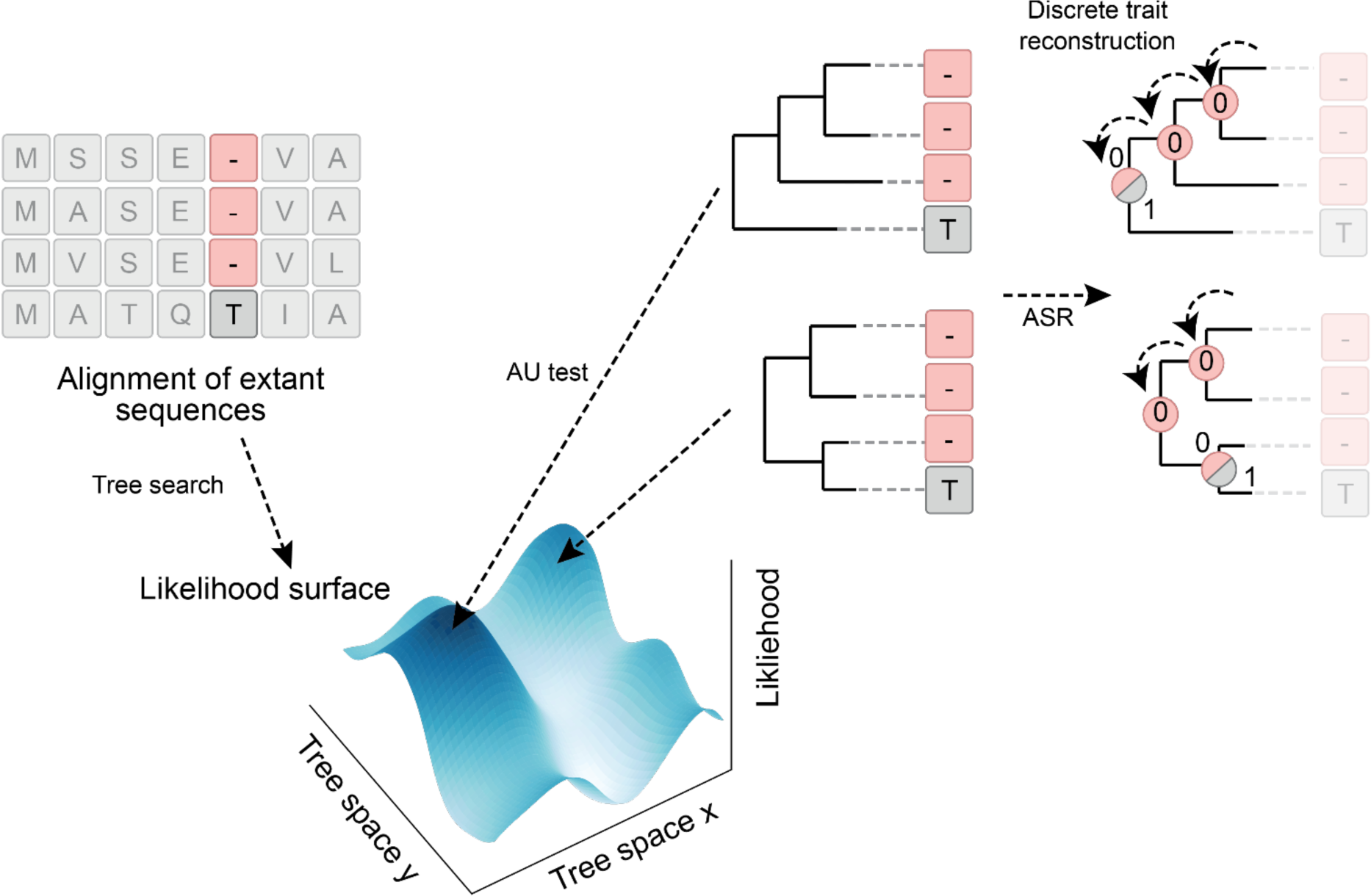
Multiplexed ASR pipeline. Extant sequences are aligned, before tree-search returns a distribution of phylogenetic trees that sample the likelihood surface over tree space. Those that fail rejection by the AU-test are used as priors for ASR. For columns in the alignment that feature gaps in ≥1% of residues, a binary value (0 or 1, representing a gap character or a non-gap character, respectively) is reconstructed by maximum likelihood in the ancestral sequence. If insertion is more likely in an ancestral sequence at a specific site than not (i.e. reconstructed likelihood ≥0.5 for node *n* at site *i*), the ancestrally reconstructed character from ASR is kept, otherwise the reconstructed character at insertion site *i* is replaced by a gap character.

**SI Fig 2.**
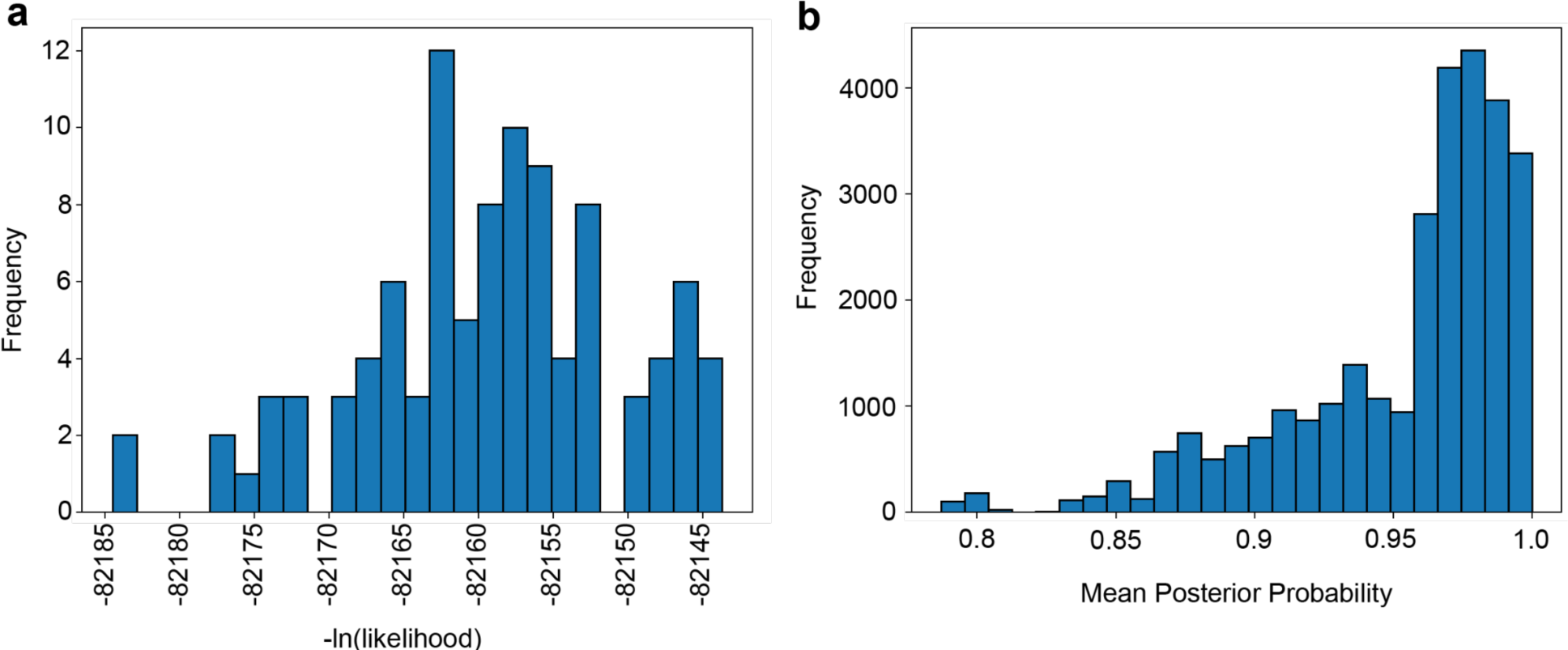
Multiplexed ASR reconstruction details. (a) -Ln(likelihood) value distribution for converged tree-searches that passed the AU test, conducted to 10000 replicates. (b) Posterior probabilities for all ancestrally reconstructed sequences.

**SI Fig 3.**
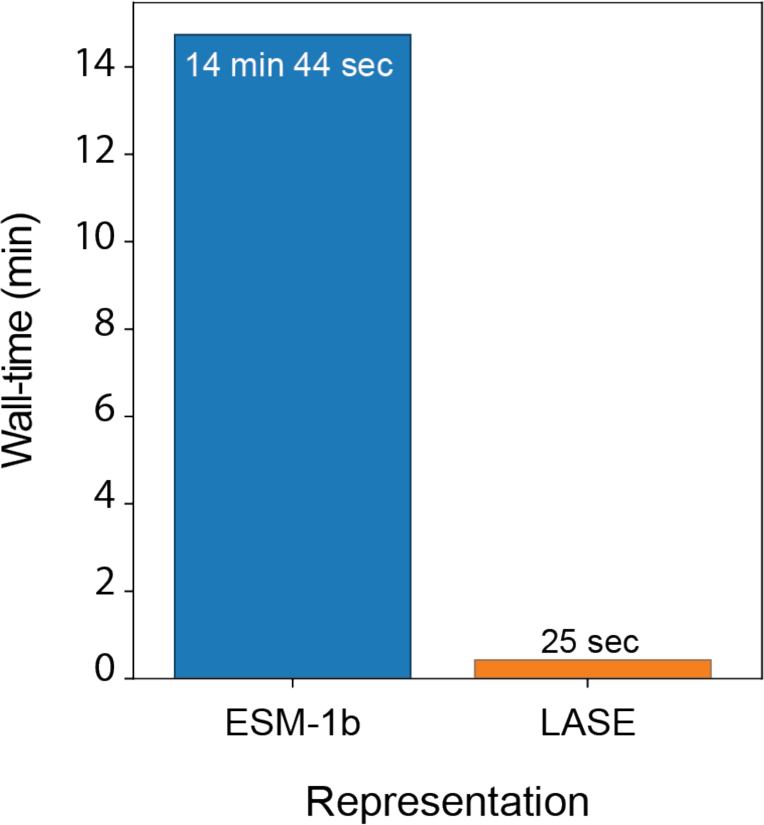
ESM-1b fine-tuning and LASE training. Both models were trained on the maximum batch size possible given GPU memory constraints. ESM-1b was fine-tuned to update all parameters on a single Nvidia A6000 (48 GB VRAM) with a batch size of 16 for a single epoch. The LASE model was trained on a single Nvidia A10 (22 GB VRAM) with a batch size of 128 for a single epoch.

**SI Fig 4.**
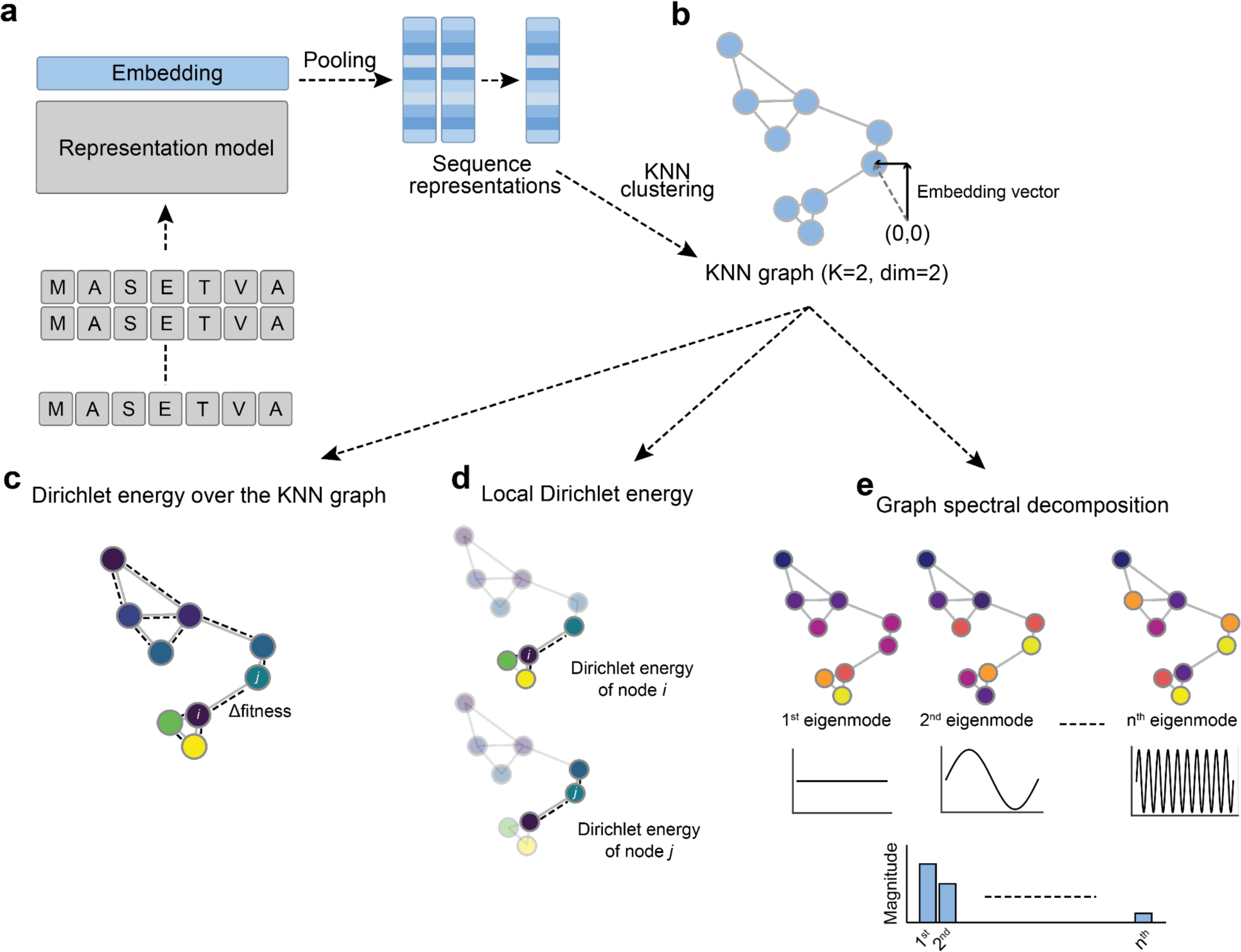
Determination of ruggedness in embedded KNN graphs. (a) Sequence embedding in the representation model. (b) Embedding vector clustering by KNN. Nodes are placed at the embedding vectors for the sequences that they represent. Edges connect each node to its *k* closest neighbors. The KNN is made undirected by making the adjacency matrix symmetrical for the graph Fourier transform to be defined. (c) Dirichlet energy is determined as the normalized sum of squared differences in fitness over all edges and is measured as a per-graph quantity. (d) The local Dirichlet energy is determined as the Dirichlet energy over subgraphs of each node and its immediate neighbors in the KNN graph, repeated for all nodes in the graph. The local Dirichlet energy is a node-wise quantity that indicates the “strained” a node is with respect to fitness, given its immediate neighbors. (e) Graph spectral decomposition uses the graph Fourier transform to determine the contribution of low order (non-complex) eigenmodes towards the fitness as a signal over the graph. It is a graph-wise quantity that relates the graph topology (the eigenvectors of the graph Laplacian) to the fitness over the graph.

### SI Text 1. Conversion of relative fitness values to catalytic efficiency

Experimentally, relative fitness is measured to be the initial rate of reaction at a given substrate concentration. In previous experiments, a 2NH concentration of 200 μM was used.

Hence, the relative rate of reaction (*v*) is proportional to 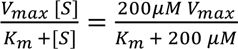.

With *k_cat_* = *V_max_*/[*E_T_*], we obtain the theoretical rate:

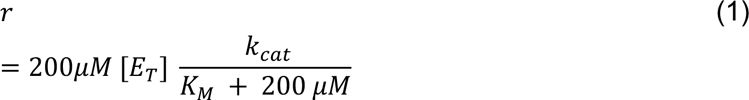

In converting *r* to the catalytic efficiency we use the below equation where α is determined based on shared data points.

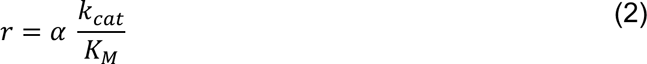

Equation (2) resembles (1) if the alpha coefficient equals the ration of the substrate by the enzyme concentration. However, the denominator of (1) has an additional term (substrate concentration) which cannot be accounted for by (2). To justify the use of (2), we calculated the theoretical (1) and calculated (2) catalytic efficiency (**SI Figure 5**) for data points for which we know both k_cat_ and K_M_ and could see that the trend is still linear (indicating use of (2) is appropriate. We also considered which k_cat_ and K_M_ values would lead to notable differences between the two equations over the range of the k_cat_ and K_M_ values we see in the dataset (**SI Figure 2b**). We can see that differences due to the use of (2) are only notable at very low K_M_ values;over the majority of the space, differences are negligible.

**SI Fig 5.**
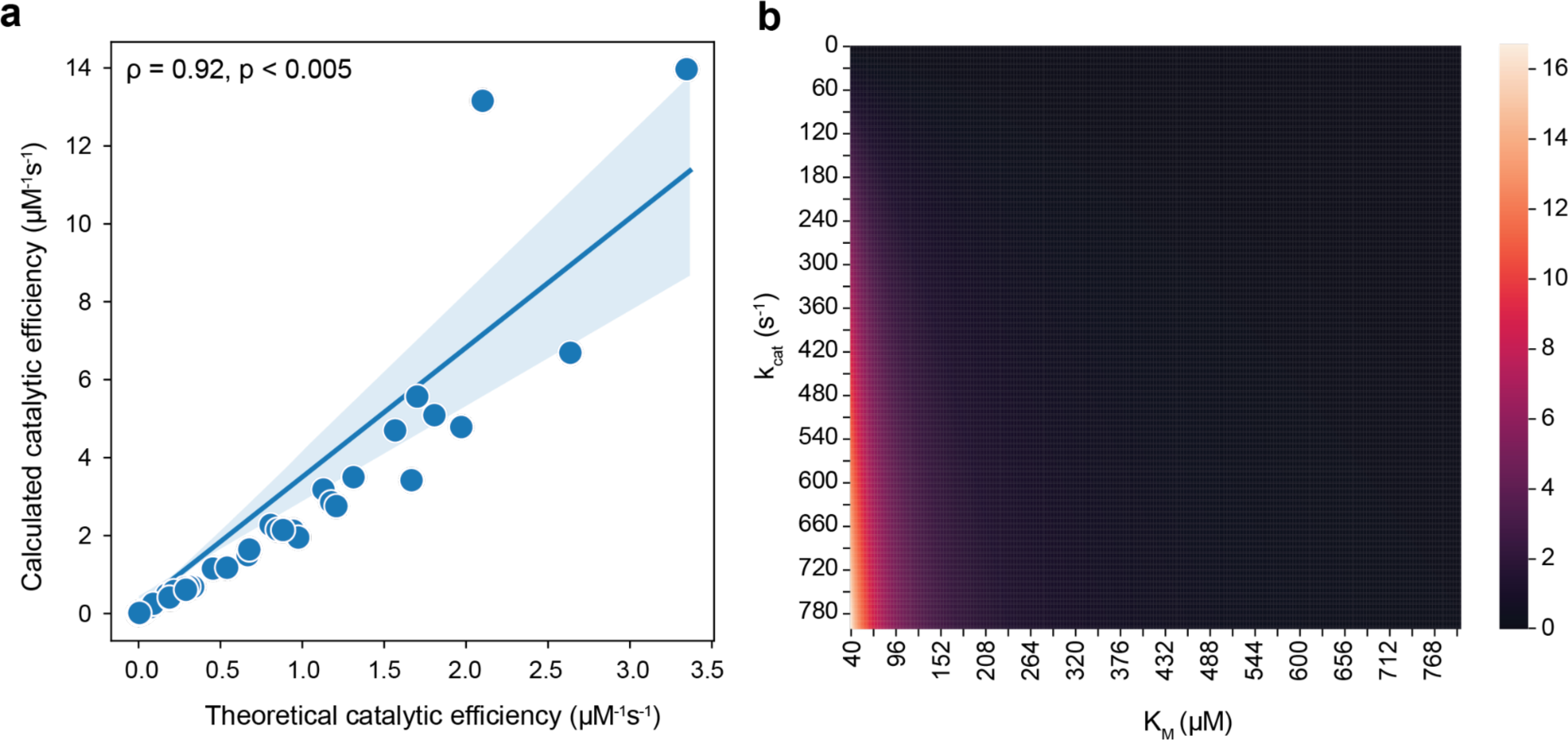
Assessment of error in converting rate to catalytic efficiency. a) Relationship between theoretical and calculated catalytic efficiency. b) Heatmap showing discrepancies between the theoretical catalytic efficiency and calculated catalytic efficiency at different k_cat_ and K_M_ values. Calculated and theoretical catalytic efficiencies in the PTE dataset share a linear relationship and deviate from each other only at low values of *K_M_*.

**SI Table 1.**
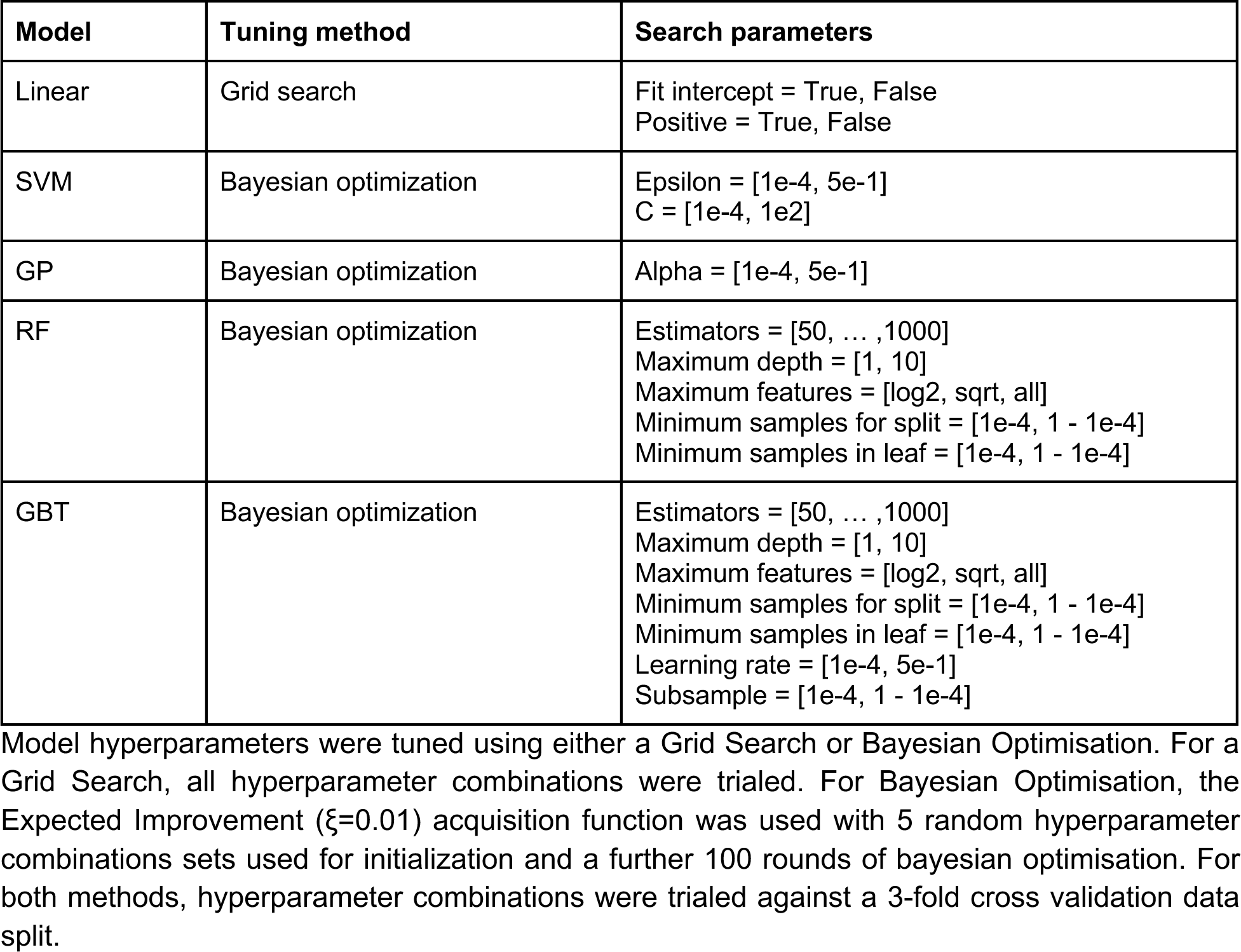
Tuning method and hyperparameter search space for regressor model optimization.

**SI Table 2.** Representation schemes used and details on extraction.

| Representation scheme | Dimensions | Details |
| --- | --- | --- |
| One-hot | $20 \times N$ | Individual amino acid vectors were concatenated length-wise for each sequence. |
| Z-Scales | $5 \times N$ | Individual amino acid vectors were concatenated length-wise for each sequence. |
| ProtFP | $8 \times N$ | Individual amino acid vectors were concatenated length-wise for each sequence. |
| Georgiev | $19 \times N$ | Individual amino acid vectors were concatenated length-wise for each sequence. |
| UniRep | 1900 | Extracted using the Jax-UniRep package. |
| ProtTrans | 1024 | Extracted using the HuggingFace feature extraction pipeline on the 'Rostlab/prot_bert_bfd' model from the HuggingFace Model Hub.<br>Special tokens were removed from the $(N + 2, 1024)$ representation to produce a $(N, 1024)$ representation which was reduced to a 1024 long representation through mean pooling. |
| ESM-1b | 1280 | Extracted using the fair-esm package with 'esm1b_t33_650M_UR50S'.<br>Special tokens were removed from the $(N + 2, 1280)$ representation to produce a $(N, 1280)$ representation which was reduced to a 1280 long representation through mean pooling. |
| ESM-2 | 1280 | Extracted using the fair-esm package with 'esm2_t33_650M_UR50D'.<br>Special tokens were removed from the $(N + 2, 1280)$ representation to produce a $(N, 1280)$ representation which was reduced to a 1280 long representation through mean pooling. |

Both amino acid descriptor sets as well as protein language models were used to embed sequences. $N = \text{Sequence length}$ .

**SI Fig 6.**
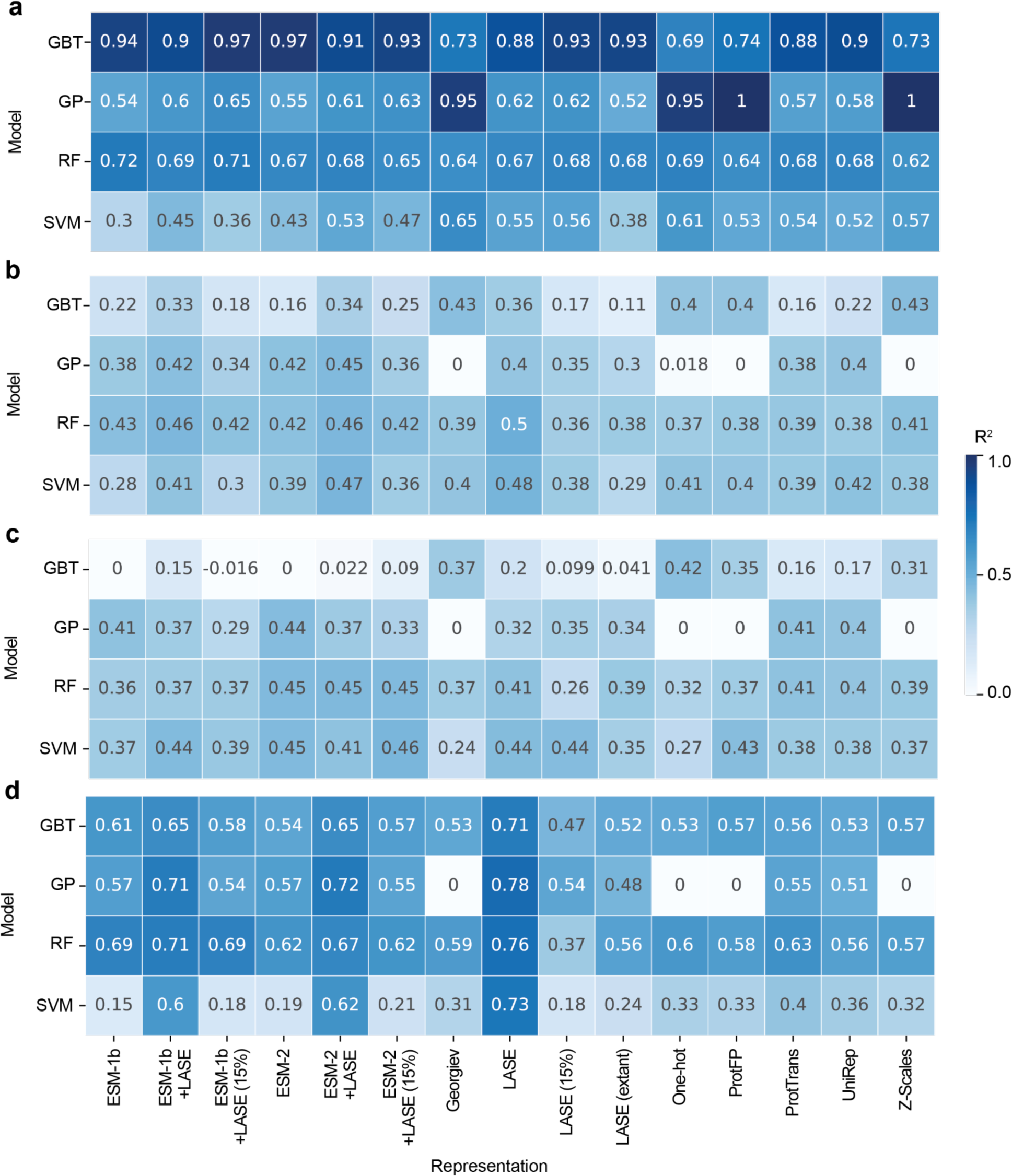
Model performance on different data splits. The R^2^ was determined for the a) training data split, b) validation data split and c) holdout data split over 10 random 20% holdout data splits with 3-fold validation for hyperparameter tuning. The final model was then trained and optimized on all available data (3-fold hyperparameter tuning on all data) before being tested on the d) interpolation data set.

